# Illuminating microbial species-specific effects on organic matter remineralization in marine sediments

**DOI:** 10.1101/705087

**Authors:** Nagissa Mahmoudi, Tim N. Enke, Steven R. Beaupré, Andreas P. Teske, Otto X. Cordero, Ann Pearson

## Abstract

Marine microorganisms play a fundamental role in the global carbon cycle by mediating the sequestration of organic matter in ocean waters and sediments. A better understanding of how biological factors, such as microbial community composition, influence the lability and fate of organic matter is needed. Here, we explored the extent to which organic matter remineralization is influenced by species-specific metabolic capabilities. We carried out aerobic time-series incubations of Guaymas basin sediments to quantify the dynamics of carbon utilization by two different heterotrophic marine isolates. Continuous measurement of respiratory CO_2_ production and its carbon isotopic compositions (^13^C and ^14^C) shows species-specific differences in the rate, quantity, and type of organic matter remineralized. Each species was incubated with hydrothermally-influenced vs. unimpacted sediments, resulting in a ~3-fold difference in respiratory CO_2_ yield across the experiments. Genomic analysis indicated that the observed carbon utilization patterns may be attributed in part to the number of gene copies encoding for extracellular hydrolytic enzymes. Our results demonstrate that the lability and remineralization of organic matter in marine environments is not only a function of chemical composition and/or environmental conditions, but also a function of the microorganisms that are present and active.

## Introduction

Microbes are an integral part of the marine carbon cycle and play a fundamental role in mediating the long-term sequestration of organic carbon in ocean waters and sediments. Through their metabolism and growth, heterotrophic microbes are key players in the transformation and remineralization of organic matter to carbon dioxide (CO_2_). Factors governing the fate of organic carbon in the ocean – either remineralization to CO_2_ or sequestration – remain underexplained. It is not clear why certain compounds or pools of organic matter are readily remineralized by heterotrophic microbes while others remain stable for millennial time scales (Middelburg *et al.*, 1993; Arndt *et al.*, 2013). The lability or availability of organic matter is dependent on intrinsic chemical and physical factors such as the molecular size and structure of the compounds as well as mineral content of the matrix (Hedges *et al.*, 2001; Rothman and Forney, 2007; Schmidt *et al.*, 2011; Lalonde *et al.*, 2012; Estes *et al.*, 2019; Hemingway *et al.*, 2019). However, it is becoming increasingly recognized that the lability and transformation of organic matter is also influenced by biological factors such as microbial community composition (Carlson *et al.*, 2004; Glassman *et al.*, 2018). Given that the lability of organic matter appears to be context-dependent, it is difficult to disentangle the interplay between these factors and, thus, elucidate the extent to which the taxonomic and functional distribution of heterotrophic microbes influences the fate of carbon in the ocean.

Although there is a need for more experimental studies resolving the relationships between the lability of organic matter and the metabolic capabilities of marine microorganisms, these interactions are difficult to assess using traditional microbiological and/or geochemical approaches. Organic matter is one of the most complex mixtures on Earth (Hedges *et al.*, 2000), and microbes derive organic carbon from varied pools that are heterogeneously distributed in space and time. Natural abundance stable carbon (^13^C) and radiocarbon (^14^C) isotopic analyses have become powerful tools for deciphering the sources and ages of natural organic matter consumed by microbes in complex environments (e.g., Petsch *et al.*, 2001; Pearson *et al.*, 2008; Mahmoudi *et al.*, 2013a, —2013b; Whaley-Martin *et al.*, 2016). Heterotrophic microorganisms carry the same Δ^14^C signatures and nearly the same δ^13^C signatures as their carbon sources; therefore, the ^13^C and ^14^C signatures of both microbial cellular components (e.g., membrane lipids) and respired CO_2_ can be used to infer microbial utilization of isotopically-distinct carbon sources (Hayes, 2001). For example, Δ^14^C measurements of bacterial lipids extracted from *Beggiatoa* mats in Guaymas Basin (Gulf of California), showed that the bacterial community was net heterotrophically consuming hydrothermal petroleum-derived carbon (Pearson *et al.*, 2005) rather than autotrophically fixing carbon from dissolved inorganic carbon (DIC) as had been suspected. More recently, δ^13^C and Δ^14^C measurements of CO_2_ collected during laboratory incubations of coastal sediment revealed that organic matter was degraded in a sequential, step-wise manner by heterotrophic bacteria: small, phytoplankton-derived compounds were degraded first, followed by petroleum-derived exogenous pollutants, and finally by polymeric plant material (Mahmoudi *et al.*, 2017).

Here, we explore the extent to which the lability, and ultimately, remineralization of organic matter is influenced by species-specific metabolic capabilities of marine microorganisms.

Using a novel bioreactor system, we carried out time-series incubations of sterilized Guaymas Basin sediments with two different heterotrophic bacterial isolates. The Isotopic Carbon Respirometer-Bioreactor (IsoCaRB) system measures the production rate and natural isotopic (Δ^14^C and δ^13^C) signature of microbially-respired CO_2_ to constrain the source and age of organic carbon that is remineralized (Beaupré *et al.*, 2016). We incubated sediments collected from on-axis (hydrothermal) and off-axis (control) sites with two different gammaproteobacteria (*Vibrio splendidus* 1A01 and *Pseudoalteromonas* sp. 3D05) representing two widely occurring genera of heterotrophic marine bacteria (Datta *et al.*, 2016; Enke *et al.*, 2018). Isotopic signatures of the respired CO_2_ showed selective utilization of different organic matter pools by these two species, while genomic analysis revealed that *Pseudoalteromonas* sp. 3D05 contained substantially more copies of genes encoding for extracellular hydrolytic enzymes, consistent with its higher observed CO_2_ production rates. These results substantiate the idea that organic matter lability and remineralization is not only a function of chemical composition and/or environmental conditions, but also a function of the microbes that are present and active in a given environment.

## Experimental Procedures

### Sampling sites

Guaymas Basin is a spreading center located in the Gulf of California with water depth of approximately 2000 m. Sediments in this basin are rich in organic carbon due to highly productive overlying waters and fast sedimentation rates (1-2 mm/yr). On-axis regions of the basin experience magmatic heating which transforms buried sedimentary organic matter into petroleum hydrocarbons that are estimated to have an average radiocarbon age of ~5000 years (Peter *et al.*, 1991; Simoneit and Kvenvolden, 1994). Hydrothermal alteration generates a complex mixture of organic compounds that permeate upward to the near-surface sediments, where it supports diverse and abundant microbial life, including microbial mats (Amend and Teske, 2005; Teske *et al.*, 2014). The total pool of sedimentary organic matter in Guaymas Basin thus encompasses a wide range of potential microbial carbon sources with distinct isotopic signatures (Pearson *et al.*, 2005), including petroleum hydrocarbons, surface-derived phytoplankton material and organic acids, allowing us to evaluate the lability of a spectrum of organic compounds. Sediment samples were chosen from on- and off-axis sampling locations in order to have contrasting organic matter pools. Sedimentary organic matter in on-axis areas would be expected to contain a diverse mixture of hydrothermal petroleum-derived organic compounds due to magmatic heating whereas sedimentary organic matter from off-axis areas would resemble more typical marine sediment.

### Sediment collection and preparation

Sediment cores were retrieved during dives with the research submersible Alvin (Woods Hole Oceanographic Institution) on a cruise to the Guaymas Basin in December 2016. Sediment cores were collected from on-axis and off-axis sites (Table S2). On-axis hydrothermal sediment cores (core 6 and 10) were retrieved during *Alvin* dive 4871 on December 23, 2016. Temperatures for these hydrothermal sediment cores (6.25 cm diameter; 9 and 20 cm long, respectively) ranged from 3°C at the seawater interface to 60-90°C at 20 cm depth; they were strongly sulfidic and covered with yellow *Beggiatoa* mats that strongly contrasted with the olive-brown bare sediments surrounding the hydrothermal hot spot. Cool off-axis sediments were collected from the northwestern ridge flanks of Guaymas Basin during *Alvin* dive 4864 on December 15, 2016, on the periphery of the “Ringvent” seep site (Table S2). Specifically, the cores 4864-6 and 4864-7 (6.25 cm diameter; 20 and 18 cm long, respectively) were taken 200m apart and had an in-situ temperature of 3-5°C as measured by the *Alvin* Heatflow probe, matching the bottom water temperature; these cores were suboxic, lacked microbial mats and matched the general seafloor sediment appearance. The sediment cores were returned to the ship within 2–4◻h of sampling, and the sediments were subsequently transported and kept in sterile glass jars at 4◻°C until the start of the experiments.

The upper 0 – 10 cm of sediment cores were homogenized and freed of carbonate species by titration to pH 2-2.5 with 10% hydrochloric acid (HCl) in an ice bath (Beaupré *et al*., 2016). Sediments were subsequently freeze dried and sterilized by gamma-irradiation via a ^137^Cs (radioactive cesium) source to receive a total dose ~40 kGy. Gamma-irradiation has been shown to minimize changes to the structure and composition of natural organic matter compared to methods that involve exposure to high temperature and pressure (e.g., autoclaving) (Berns *et al.*, 2008). Sediments were confirmed to be sterilized as follows: First, sterilized sediment was diluted in Tibbles-Rawlings (T-R) minimal medium (see supplemental section for detailed recipe) and plated onto Marine Broth 2216 (Difco #279110) 1.5% agar (BD #214010) plates to assess the growth of any colonies. Second, since spores frequently grow upon re-wetting (Kieft, 1987), several grams of sterilized sediment were incubated in ~50 ml of in T-R minimal medium for 5 days on a benchtop shaker and subsequently plated onto Marine Broth agar plates to observe the growth of any colonies. In both cases, no colonies were observed. Lastly, ~10 g of sterilized sediment was incubated with 1 l of T-R minimal medium in the IsoCaRB system under oxic conditions (20% O_2_) for 72 hours; the concentration of CO_2_ during this incubation remained at baseline and the total quantity of collected CO_2_ was indistinguishable from the background of the system indicating that no microbial respiration was observed.

### Bacterial strains and culture conditions

Two model marine strains (*Vibrio sp.* 1A01 and *Pseudoalteromonas sp.* 3D05) were previously isolated from coastal ocean water samples (Canoe Beach, Nahant, MA, USA; 42°25′11.5′′ N, 70°54′26.0′′ W) for their ability to degrade complex substrates, specifically model chitin particles (Datta *et al.*, 2016; Enke *et al.*, 2018). The strains were inoculated from glycerol stocks into 10 ml of liquid Marine Broth 2216 (Difco #279110) in a 125 ml Erlenmeyer flask and incubated at room temperature with shaking at 150 rpm. Upon reaching log-phase (5 hours), cells were transferred (2% inoculation) into 50 ml of T-R minimal medium supplemented with 0.5% (w/v) *N*-acetylglucosamine (Sigma-Aldrich #G4875) as a carbon source and incubated ~12 hr at room temperature and 150 rpm. Cells were transferred to 50 ml of fresh T-R minimal medium supplemented with 0.5% (w/v) *N*-acetylglucosamine until they reached a desired cell density of 5 × 10^8^/ml. Cell density was monitored by measuring optical density (OD) 600 nm, based on a calibration curve between OD and colony forming units (CFUs). Once cultures had reached the desired cell density, 50 ml of culture was harvested by centrifugation for 10 min at 3000 rpm (Beckman Coulter Allegra X30-R Centrifuge) and washed two times with T-R minimal medium. The cell pellet was then resuspended in 1 ml of T-R minimal medium and injected into the IsoCaRB system using a 3 ml syringe (BD Biosciences # 309657) and a 20-gauge needle (Sigma-Aldrich #Z192511).

### Incubations in the IsoCaRB system

For each experiment, approximately 22 g (dry weight) of decarbonated, sterilized sediment was incubated at room temperature (~22°C) in a custom Pyrex vessel containing 2 l of T-R minimal medium. The slurry was continuously stirred (90 rpm) under aerobic conditions to provide an unlimited supply of O_2_. The sediment slurry was subsampled every 12 hours to track the number of viable cells. Approximately 1 ml of sediment slurry was serially diluted and 100 μL was plated in 10^−4^ dilution in triplicate on MB2216 agar plates using rattler beads (Zymo #S1001); colony forming units (CFU) were counted after 72 hours.

Details regarding the standard operating procedure for the IsoCaRB system, including sterilization and assembly, sample preparation, and CO_2_ collection and purification are described in Beaupré *et al.*, (2016). Briefly, the system is first purged of residual atmospheric CO_2_ by sparging with CO_2_-free helium for 48 hours. Gas flow is then changed to 100 ml min^−1^ of 20% O_2_ in helium approximately one hour prior to injection of the bacterial cells. Respiratory CO_2_ is carried to an online infrared CO_2_ analyzer (Sable Systems CA-10) where concentrations are quantified in real time and continuously logged to a desktop PC using a custom LabVIEW program (National Instruments). This CO_2_ is continuously collected as successive fractions in custom molecular sieve traps and CO_2_ is recovered from the traps by baking (600°C for 75 min) under vacuum within ~24 hours of collection, cryogenically purified, quantified, and flame-sealed in Pyrex tubes. Each experiment was allowed to proceed until CO_2_ concentrations reached near-baseline values.

Gaseous CO_2_ concentration measurements were corrected for baseline drifts and then rescaled to agree with the higher-precision manometric yields obtained from the trapped CO_2_ (Beaupré *et al*., 2016). These normalized CO_2_ concentrations were corrected for the confounding effects of mixing in the culture vessel headspace and decreasing slurry volume (as described in Mahmoudi *et al.*, 2017) to calculate the rate of CO_2_ generation per unit volume of growth medium (μg C l^−1^ min^−1^), which serves as a proxy for the microbial CO_2_ production rate.

A total of six incubations were performed: one on-axis and two off-axis samples for each of the two bacterial isolates. The rate of microbial CO_2_ production and CFUs/ml between the replicate incubations of off-axis sediments were highly reproducible for both isolates, and the total quantity of respired CO_2_ collected varied by less than 0.1 mg between replicate incubations (Figures S2). Thus, CO_2_ fractions from a single off-axis incubation for each isolate were submitted for δ^13^C and Δ^14^C to minimize costs.

### Isotopic analysis of CO_2_

CO_2_ fractions were sent to the National Ocean Sciences Accelerator Mass Spectrometry (NOSAMS) Facility at the Woods Hole Oceanographic Institution. An aliquot of each sample was split for ^13^C measurement, with the remainder reduced to graphite (Vogel *et al*., 1987) for ^14^C measurement by accelerator mass spectrometry (AMS). All isotopic data are corrected for background contamination associated with the IsoCaRB system (e.g., sparging gases) and the T-R minimal medium (total ~35 μg C/day, δ^13^C = 1.7‰, Δ^14^C = −141‰) as described in Mahmoudi *et al.*, (2017). Stable isotope values (δ^13^C) are reported vs. the VPDB standard. Radiocarbon values (^14^C) are reported in Δ^14^C notation, where Δ^14^C is the relative deviation from the ^14^C/^12^C ratio of the atmosphere in 1950 (Stuiver and Polach, 1977).

### Genomic characteristics

The genomes of *Vibrio* sp. 1A01 and *Pseudoalteromonas* sp. 3D05 were previously sequenced, assembled and deposited in project PRJNA478695 by Enke *et al.*, (2018) (Table S3). The remineralization of organic matter in marine environments is thought to be largely dependent on the production of extracellular hydrolytic enzymes which initiate remineralization by hydrolyzing substrates to sufficiently small sizes in order to be transported across cell membranes (Arnosti, 2011). The copy number of genes encoding for extracellular hydrolytic enzymes was quantified by searching for putative genes in the annotated genomes for known classes of hydrolytic enzymes using GENEIOUS PROv7.1. Subsequently, identified genes were translated and assessed using SignalIP (v 5.0) to determine whether they contained secretory signal peptides needed for extracellular transport across the cell membrane (Nielsen *et al.*, 2007; Armenteros *et al.*, 2019). Lastly, any genes involved in the hydrolysis of polysaccharides were also searched in the CAZy database (Cantarel *et al.*, 2008) to confirm they matched homologues of known hydrolytic enzymes.

## Results

### Microbial CO_2_ production and carbon utilization

Incubation of sterilized on- and off-axis Guaymas Basin sediment revealed a similar respiration pattern for both species. CO_2_ production rates rapidly increased with the onset of the incubation and peaked within the first 12 to 20 hours, gradually decreasing to near baseline values within ~4 days for the *Vibrio* sp. 1A01 and ~6 days for the *Pseudoalteromonas* sp. 3D05 (Figure 1; Figure 2). Likewise, cell growth was observed to increase within the first 12 to 20 hours, peaking at 1.2 – 2.1 × 10^7^ CFU/ml (Figure S1). Following this initial growth phase, cell densities remained steady at ~ 1 × 10^7^ CFUs/ml and slightly decreased toward the end of the incubation (Figure S1; Table S4).

**Figure 1.**
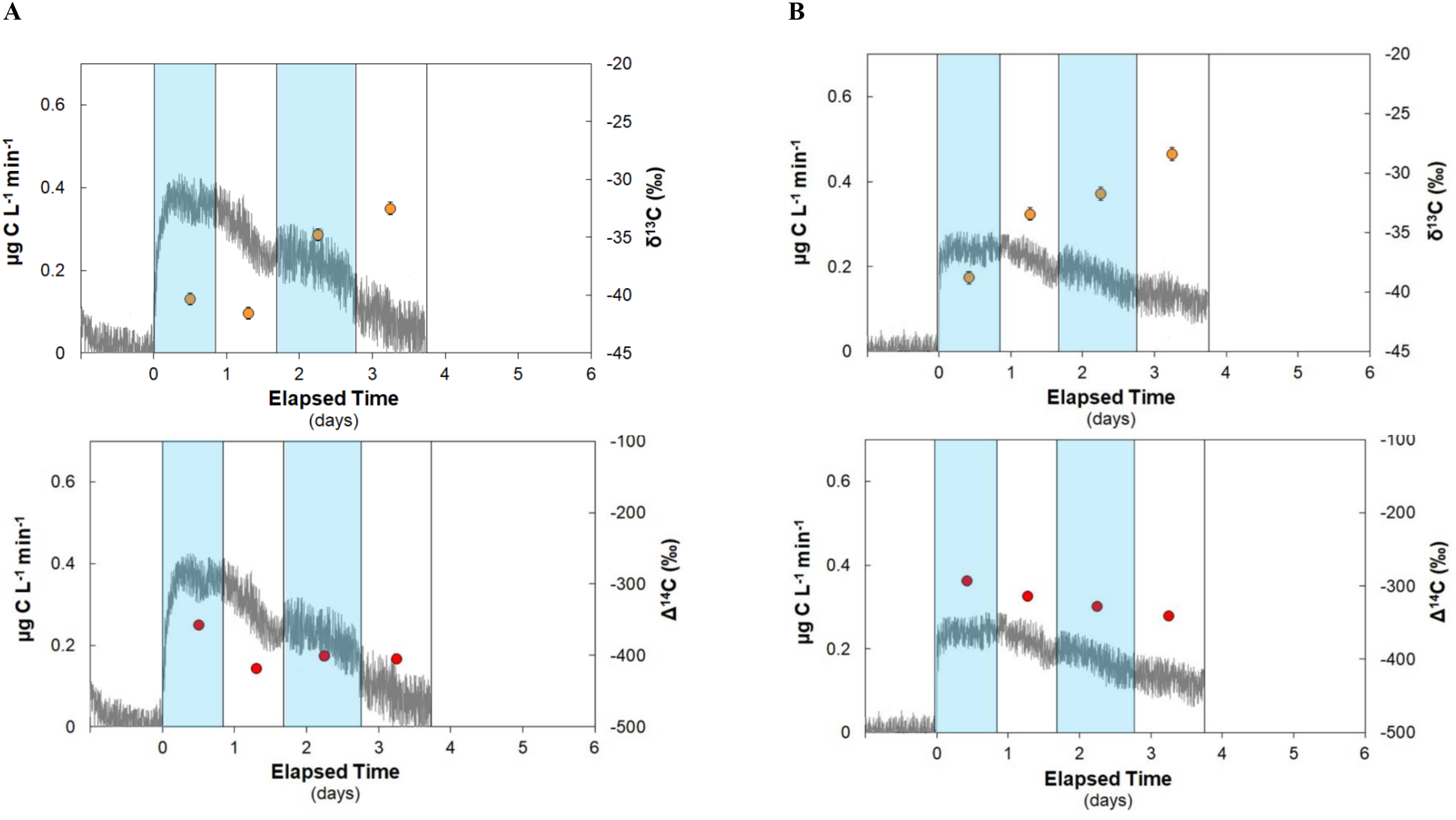
Microbial CO_2_ production rates (gray line), δ^13^C and Δ^14^C (circles) signatures of respired CO_2_ observed during incubation of *Vibrio* sp. 1A01 with (A) on-axis and (B) off-axis Guaymas Basin sediment. The width of each box spans the time interval during which each CO_2_ fraction was collected for isotopic analysis, with the corresponding data point plotted at the mid-point for each fraction.

**Figure 2.**
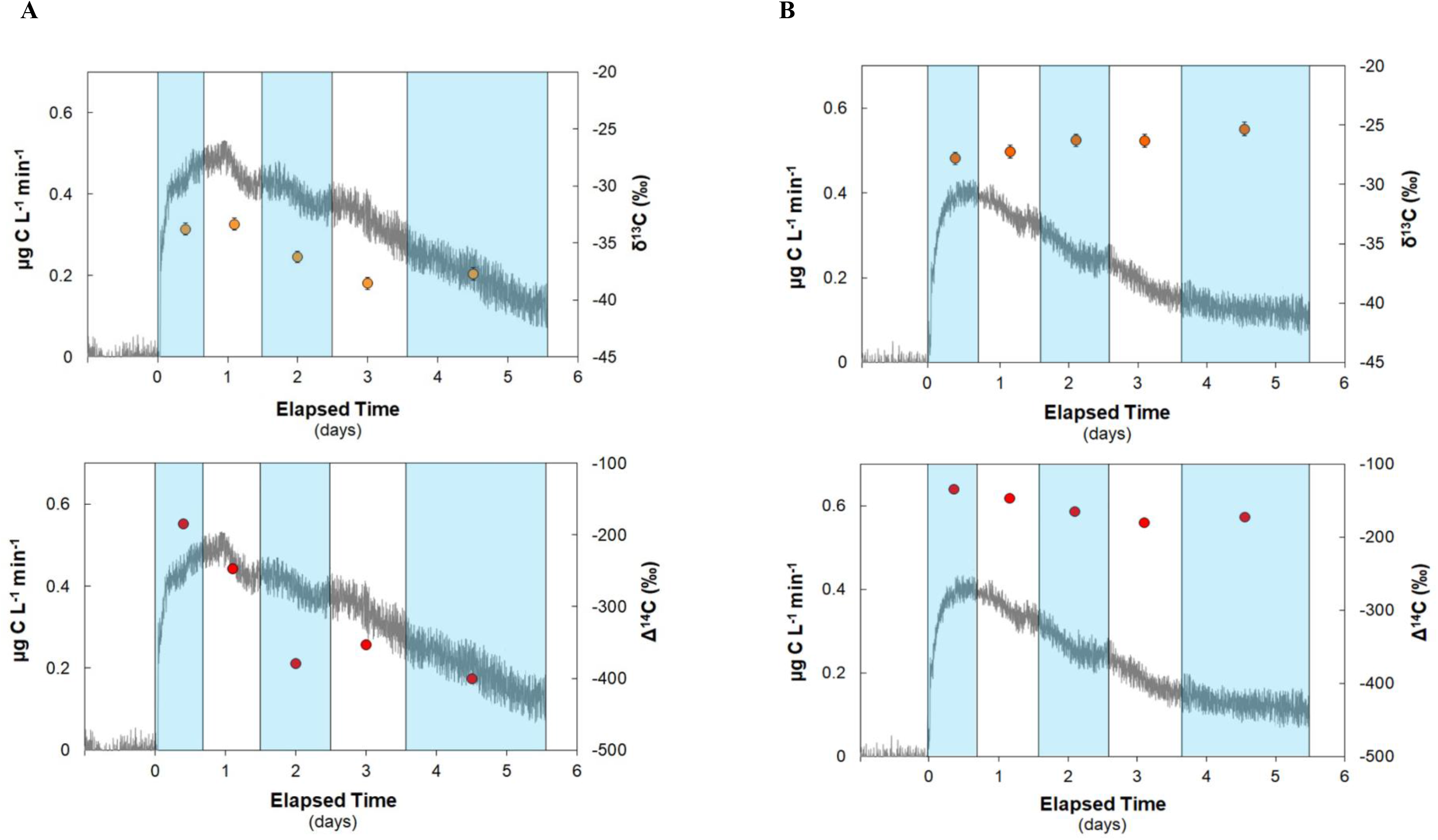
Microbial CO_2_ production rates (gray line), δ^13^C and Δ^14^C (circles) signatures of respired CO_2_ observed during incubation of *Pseudoalteromonas* sp. 3D05 with (A) on-axis and (B) off-axis Guaymas Basin sediment. The width of each box spans the time interval during which each CO_2_ fraction was collected for isotopic analysis, with the corresponding data point plotted at the mid-point for each fraction.

Higher rates of CO_2_ production were observed with *Pseudoalteromonas* sp. 3D05 compared to *Vibrio* sp. 1A01 in all cases. The maximum rates of CO_2_ production for *Vibrio* sp. 1A01 were 0.4 μg C L^−1^ min^−1^ during incubation with the on-axis sediment and 0.3 μg C L^−1^ min^−1^ during incubation with the off-axis sediment (Figure 1), whereas *Pseudoalteromonas* sp. 3D05 had a maximum rate of 0.5 μg C L^−1^ min^−1^ and 0.4 μg C L^−1^ min^−1^ (Figure 2), respectively, for the equivalent incubations. Interestingly, CO_2_ production rates were higher for both species during incubation with on-axis sediment, rather than the non-hydrothermally active off-axis sediment. This is consistent with previous work showing that hydrothermal petroleum-derived carbon in Guaymas Basin is labile and readily utilized by *in situ* microorganisms (Pearson et al., 2005).

To determine the source and age of organic matter being remineralized by each species, successive CO_2_ fractions were collected for natural abundance isotopic analysis (δ^13^C and Δ^14^C). A total of four CO_2_ fractions were collected for the *Vibrio* sp. 1A01 incubations and five CO_2_ fractions for *Pseudoalteromonas* sp. 3D05 incubations (Figure 4; Table S5). The duration of each fraction ranged from 15 to 48 hours to ensure that each fraction contained ≥0.5 mg of carbon. A total of 2.3 mg and 1.7 mg of respired carbon was collected during incubation of *Vibrio* sp. 1A01 with on- and off-axis sediment, respectively. Approximately twice as much respired carbon was collected during incubation of *Pseudoalteromonas* sp. 3D05, with 4.6 mg and 3.3 mg collected for on- and off-axis sediment.

Temporal trends in respired CO_2_ δ^13^C values differed between incubations with *Vibrio* sp. 1A01 and *Pseudoalteromonas sp. 3D05*, confirming differential utilization of organic matter by each species. For *Vibrio* sp. 1A01, δ^13^C signatures of CO_2_ fractions became more ^13^C-enriched over time, ranging from −40 to −33‰ and −39 to −28‰ for on- and off-axis sediment, respectively (Figure 1; Table S5). In contrast, δ^13^C signatures of respired CO_2_ fractions became ^13^C-depleted (−34 to −38‰) during incubation of *Pseudoalteromonas* sp. 3D05 with on-axis sediment but ^13^C-enriched (−28 to −25‰) during incubation with off-axis sediment (Figure 2; Table S5).

The Δ^14^C signatures of respired CO_2_ decreased with time during all incubations, indicative of utilization of progressively older compounds. The Δ^14^C signatures of respired CO_2_ fractions of *Vibrio* sp. 1A01 ranged from −357 to −405‰ and −292 to −341‰ during incubation with on- and off-axis sediment (Figure 1; Table S5). For *Pseudoalteromonas* sp. 3D05, Δ^14^C signatures of respired CO_2_ fractions ranged from −185 to −400‰ and −134 to −172‰ during incubation with on- and off-axis sediment (Figure 2; Table S5). The Δ^14^C signatures of respired CO_2_ fractions of *Pseudoalteromonas* sp. 3D05 were generally more positive (by ~150‰) than the equivalent fractions of *Vibrio* sp. 1A01, indicating utilization of a greater proportion of recently photosynthesized carbon such as phytoplankton-derived material. Both species had lower Δ^14^C signatures during incubation with on-axis sediment, consistent with the utilization of hydrothermal petroleum-derived carbon (Δ^14^C = −431 ± 55‰; Simoneit and Kvendvolden, 1994).

### Extracellular hydrolytic enzymes

The genomes of both species contain multiple copies of genes encoding for chitinases and peptidases which are involved in the degradation of chitin and proteins, respectively (Figure 3). Both species have a comparable number of gene copies for chitinases, specifically, *Vibrio* sp. 1A01 had 5 gene copies and *Pseudoalteromonas* sp. 3D05 had 7 gene copies. However, *Pseudoalteromonas* sp. 3D05 has substantially more gene copies for extracellular peptidases with a total of 30 copies whereas *Vibrio* sp. 1A01 has 7 copies. In addition, *Pseudoalteromonas* sp. 3D05 has a single gene copy encoding for a glucosidase which is involved in the degradation of complex carbohydrates while *Vibrio* sp. 1A01 did not contain any genes encoding for extracellular glucosidases.

**Figure 3.**
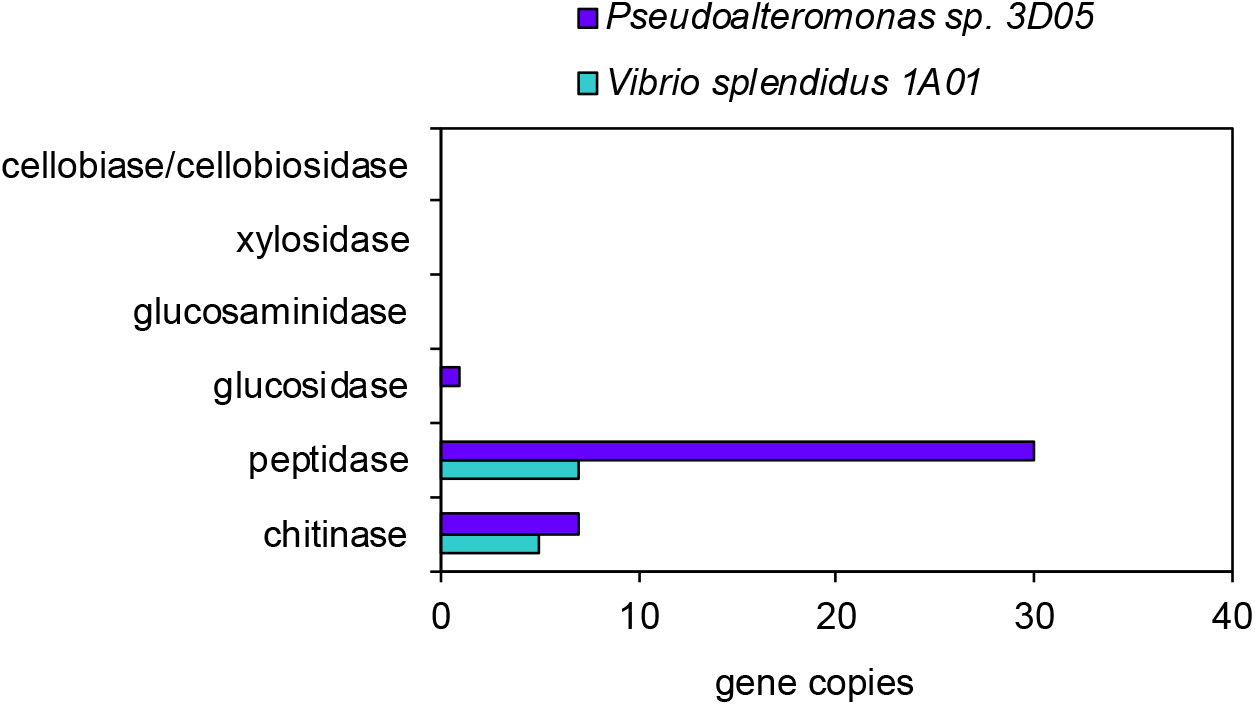
Genomic features of isolates used for incubations with Guaymas Basin sediment. The total copy number of genes encoding for known extracellular enzymes was determined from annotated genomes. Gene sequences were translated and assessed using SignalIP (v 5.0) to determine whether they contained secretory signal peptides needed for extracellular transport. Both *Vibrio* sp. 1A01 and *Pseudoalteromonas* sp. 3D05 were found to have similar copy numbers of chitinases; however, *Pseudoalteromonas* sp. 3D05 was found to have ~4 times more genes encoding for extracellular peptidases.

**Figure 4.**
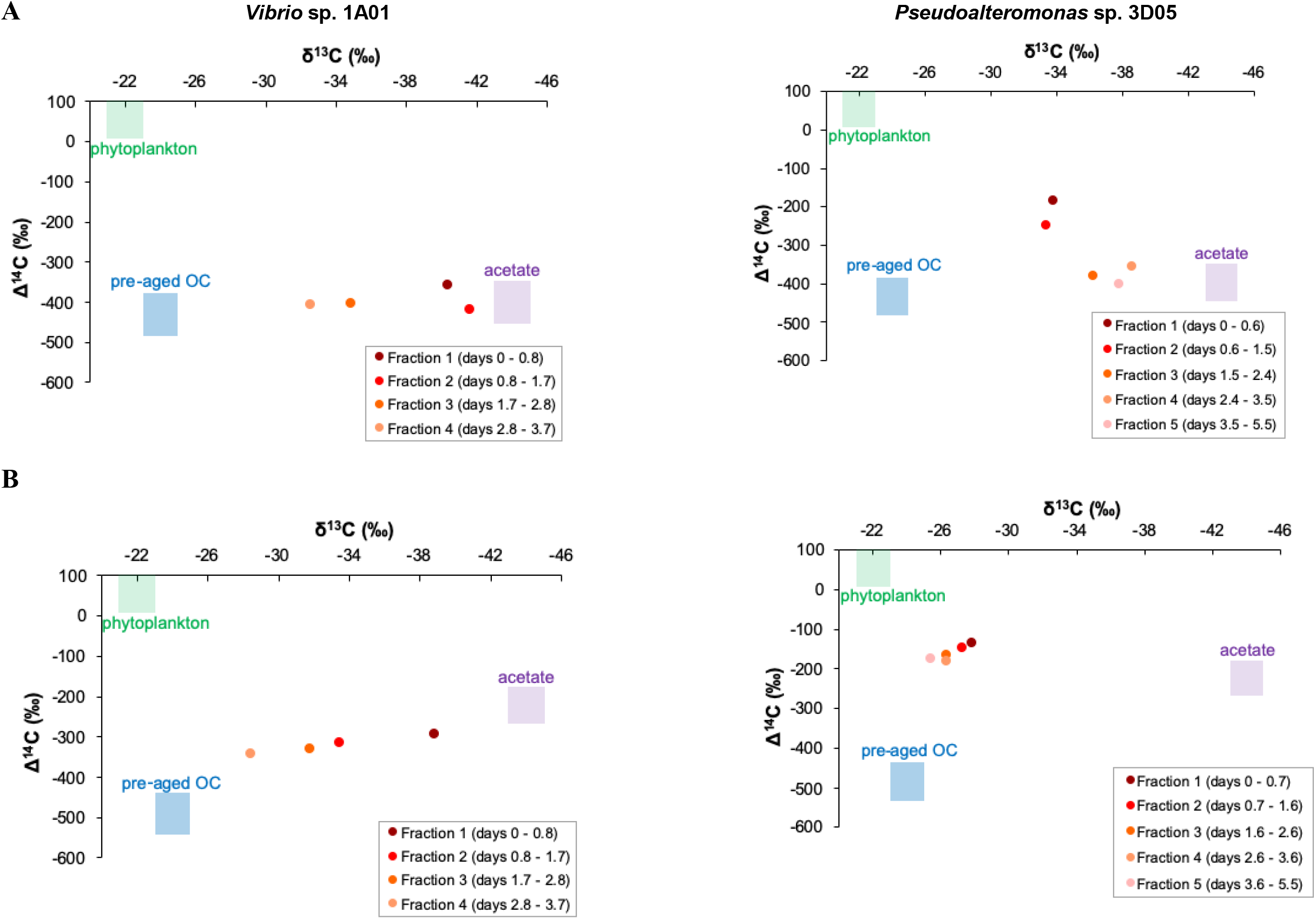
Plot of stable carbon (δ^13^C) versus radiocarbon (Δ^14^C) signatures of respired CO_2_ collected from incubation of *Vibrio* sp. 1A01 and *Pseudoalteromonas* sp. 3D05 with (A) on-axis and (B) off-axis Guaymas Basin sediment. Also shown are assumed δ^13^C values and Δ^14^C values for potential microbial carbon sources.

## Discussion

Resolving the mechanisms that control whether and/or how organic matter is consumed by microbes is crucial for understanding how carbon is processed and stored in the ocean. Previous work using bottle incubations with seasonally accumulated dissolved organic matter (DOM) and natural seawater from different depths observed differences in DOM degradation when this same carbon pool was incubated with natural seawater from contrasting depths (Carlson *et al.*, 2004, Letscher *et al.*, 2015), which suggested that microbial community structure is an important control on the remineralization of organic matter. This view is consistent with observations of striking contrasts in the abilities of water column and sedimentary microbial communities to utilize high molecular weight substrates such as polysaccharides (Arnosti 2000; 2008), contrasts that parallel differences in microbial community composition (Teske *et al.*, 2011; Cardman *et al.*, 2014). However, empirically testing the extent to which the taxonomic and/or functional characteristics of microorganisms can give rise to differences in remineralization is analytically challenging. Our study leverages a novel bioreactor system to track and quantify the dynamics of carbon utilization by two different marine isolates by continuously measuring CO_2_ production and carbon isotopes associated with the remineralization of natural organic matter (Beaupré *et al.*, 2016). Using this approach, we observed species-specific differences in the rate, quantity, and type of organic matter remineralized. Our results demonstrate that the lability and remineralization of organic matter is not exclusively affected by chemical and physical factors but that it will also be dependent on the microbes that are present and active in a given environment.

### Species-level selectivity of organic carbon substrates

In addition to observing differences in remineralization rates and yields between species, the observed trends in isotopic signatures of respired CO_2_ indicate differences in the types of carbon compounds that were degraded by each species (Figure 4), thereby influencing the types of carbon compounds left behind and the terminal carbon storage. Given the diverse pools of sedimentary organic matter available for microbial utilization, we attempted to constrain the respective contribution of multiple carbon pools to microbial respiration using a mass balance model. Sedimentary organic matter generally is believed to be dominated by phytoplankton-derived organic compounds (Wakeham *et al.*, 1997, Burdige 2007), especially in locations with fast sedimentation rates and productive overlying waters. In addition, previous measurements of organic acids in Guaymas Basin sediment have found high concentrations of acetate in porewaters (Martens, 1990), consistent with the notion that microbial acetogenesis – and/or fermentation to low-molecular-weight organic acids – is widespread in marine sediments (Lever *et al.*, 2010; Lever, 2012; He *et al.*, 2016). Marine sediments can also contain inputs of older (^14^C-depleted) carbon that can be derived from numerous sources. Aged terrigenous organic matter, derived from land plants, contributes significant organic matter to continental margin sediments (Raymond and Bauer, 2001; Schefuß *et al.*, 2016).

Because we cannot be certain of the exact source and composition of this older carbon within our sediments, we simply refer to this endmember as pre-aged organic carbon (OC) in our mass balance model. However, in the case of sediments collected from on-axis sampling locations, we assume that the pre-aged OC endmember is dominated by hydrothermal-derived petroleum compounds. Guaymas Basin is relatively unusual in that it experiences magmatic heating which helps mobilize pre-aged carbon through the generation of hydrothermal petroleum (Peter *et al.*, 1991). This petroleum eventually migrates vertically and discharges at the sediment water interface where it is readily used by bacterial communities (Pearson *et al.*, 2005). Previous studies have reported that surface sediments in on-axis areas are rich in petroleum hydrocarbons with concentrations comparable to those in highly polluted and industrial areas (Kawka and Simoneit, 1990; Simoneit and Lonsdale, 1982). Thus, our mass balance model presumed three major carbon sources: phytoplankton-derived compounds, microbially-produced acetate and/or other fermentation products, and pre-aged OC.

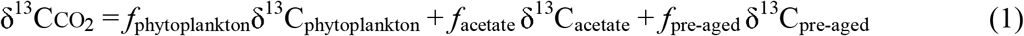

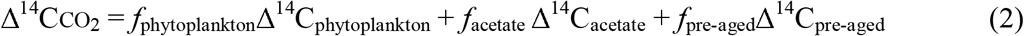

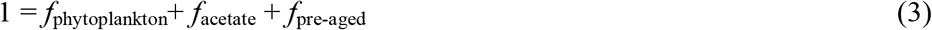

where δ^13^C_CO_2__ and Δ^14^C_CO_2__ correspond to the measured isotopic values of the respired CO_2_. The phytoplankton endmember was assumed to have a commonly accepted δ^13^C value of −22 ± 1‰ (Coleman and Fry, 1991) and the Δ^14^C was set to −12 ± 50‰, after accounting for the input of “bomb carbon” derived from atmospheric testing of nuclear weapons in the 1950s and 1960s. For on-axis sediment incubations, the pre-aged OC endmember was set to the previously measured values of Guaymas petroleum oils (δ^13^C = −23 ± 1‰, Δ^14^C = −431‰ ± 55‰, Simoneit and Schoell, 1995; Simoneit and Kvenvolden, 1994). For off-axis sediment incubations, the δ^13^C value of pre-aged OC endmember was set to −24 ± 1‰, based on previous measurements of organic carbon in the Colorado River (δ^13^C = −20 to −27‰; Daessle *et al.*, 2017) which feeds directly into the Gulf of California, while the Δ^14^C value was set to −490 ± 50‰ after iteratively constraining potential ranges for this endmember. Lastly, for the acetate endmember, the δ^13^C value was set to −44 ± 1‰ for both on- and off-axis sediment incubations, based on the expected fractionation heterotrophic acetogenesis which would have resulted in acetate as being ~19.5‰ more depleted relative to initial substrate (Freude and Blaser, 2016). Most of the acetate in marine sediments appears to be generated through heterotrophic rather than autotrophic acetogenesis (Heuer *et al.*, 2009; Lever *et al.*, 2010; Ijiri *et al*., 2010), consistent with the idea that autotrophic acetogens would be easily outcompeted by other microbes (e.g., methanogens) in marine sediments due to the very low energy yield of this metabolism (Lever, 2012). For the on-axis sediment incubations we constrained the ranges of possible acetate Δ^14^C signatures and set the Δ^14^C value of acetate to −400 ± 50‰, which suggests hydrothermal-derived petroleum compounds are the likely source of substrates for fermentation. For the off-axis sediment incubations we set the Δ^14^C value of acetate to −226 ± 50‰, which reflects the measured value for the total organic carbon pool and assumed that it reflected the average material being fermented. Details on the assumptions and uncertainty associated with each endmember value can be found in the Supporting Information. Fractional contributions (*f’*s) and their uncertainties were estimated as the means and standard deviations of solutions to Equations 1 – 3, which were solved by Monte Carlo resampling (10,000 times) with normally-distributed pseudo-random noise added to each isotope measurement (Δ^14^C ± 5 ‰; δ^13^C ± 0.5 ‰) and end-member isotopic signature.

Assuming that these are the primary sources of carbon available in the sediment, we predicted the fractional utilization of all three sources for each species using this mass balance model, including which pools were preferentially respired at different times of the incubation (Figure 5). *Vibrio* sp. 1A01 was found to respire CO_2_ derived primarily from acetate and pre-aged OC. Specifically, acetate was a dominant initial carbon source, accounting for 82% and 75% of all respired CO_2_ during the first day of incubation with on- and off-axis sediment, respectively. The relative contribution of acetate to respiration decreased with time for *Vibrio* sp. 1A01 and concurrently there was increased utilization of pre-aged OC such that it accounted for 50% and 60% of all respired CO_2_ during the last day of incubation with on- and off-axis, respectively. This trend may be due to progressive consumption of the acetate pool since it was so heavily utilized early on. *Pseudoalteromonas* sp. 3D05 respired all three carbon sources with different trends observed during incubation with on- vs. off-axis sediment. During incubation with on-axis sediment, there was substantial utilization of acetate such that 50 to 77% of all respired CO_2_ was derived from this pool throughout the duration of the experiment. Similarly, pre-aged OC was utilized by *Pseudoalteromonas* sp. 3D05 over the course of the on-axis incubation; however, it remained a minimal carbon source and never accounted for more than ~30% of respired CO_2_. Interestingly, *Pseudoalteromonas* sp. 3D05 derived a substantial portion (41-56%) of all respired CO_2_ from phytoplankton-derived material early in the incubation with on-axis sediment; in contrast, there was limited utilization (<20%) of this pool by *Vibrio* sp. 1A01. This contrast was more evident during incubation with off-axis sediment where phytoplankton-derived material was the predominant carbon source for *Pseudoalteromonas* sp. 3D05, accounting for ~60% of all respired CO_2_. The remainder of its carbon was derived from a mixture of both acetate and pre-aged OC whose contributions to respired CO_2_ over time decreased and increased, respectively.

**Figure 5.**
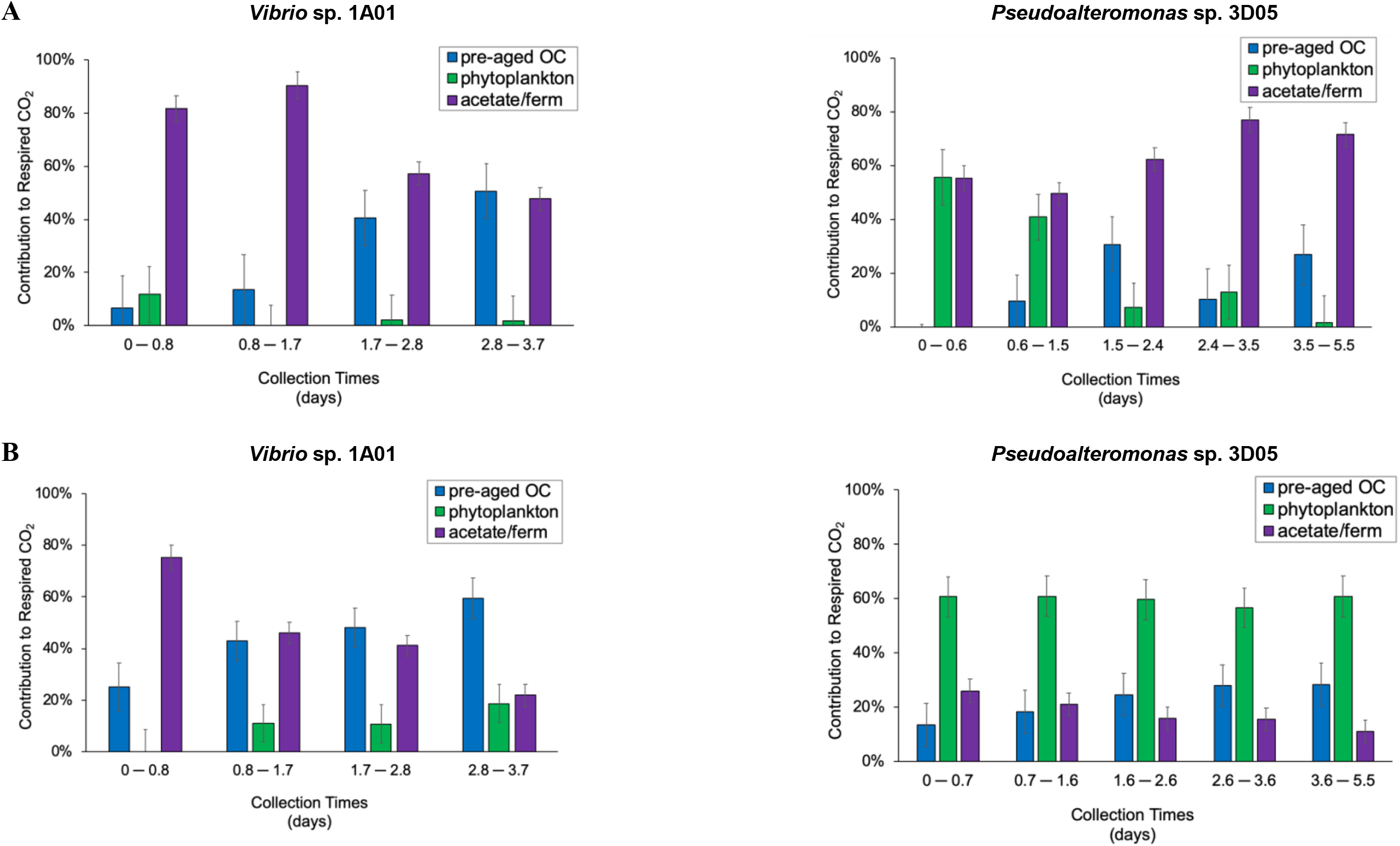
Differential utilization of carbon substrates between species. Estimated contributions of potential carbon sources to respired CO_2_ during incubations of *Vibrio* sp. 1A01 and *Pseudoalteromonas* sp. 3D05 with (A) on-axis and (B) off-axis Guaymas Basin sediment. Relative contributions were estimated using a three-end mass balance model. Percentages and uncertainties were estimated as means and standard deviations of solutions to 3 simultaneous mass-balance equations that were solved 10,000 times, in which normally-distributed pseudo-random noise was added to each isotope ratio measurement and isotopic signature.

The results of the mass balance model reveal unique trends in carbon utilization between species. Both species readily consume acetate (or other low-molecular-weight fermentation products – here referred to generically as acetate) as a carbon source. This is consistent with expectations, given that acetate is an energetically-rich compound that is labile and readily accessible to bacteria. The heavy consumption of acetate by *Vibrio sp.* 1A01 may due to the fact that it has more limited hydrolytic capabilities which may be required to access other carbon pools; in contrast, *Pseudoalteromonas* sp. 3D05 readily utilized substrates from all carbon pools particularly phytoplankton-derived carbon. Given that proteins are a major component of phytoplankton cells (25-50%; Emerson and Hedges, 2008) and phytoplankton-derived carbon pools in degraded marine sediments are believed to be protein-rich (Cowie and Hedges, 1984), it would be expected that compounds within this pool need to be degraded extracellularly before being transported into bacterial cells. Likewise, we found that *Pseudoalteromonas* sp. 3D05 contained ~4 times more genes copies for extracellular hydrolytic peptidases compared to *Vibrio* sp. 1A01, consistent with its greater fractional reliance on carbon pools comprised of larger substrates.

### Microbial controls on organic matter remineralization

While much is known about how abiotic conditions and intrinsic chemical properties may limit the remineralization of organic matter, the extent to which this process may be influenced by the microorganisms is just beginning to be explored. A greater understanding of the microbial characteristics or traits that affect remineralization will improve our ability to predict the fate of organic carbon in marine ecosystems. A large proportion of organic matter, particularly in marine sediments, is macromolecular and physically too large to be utilized directly by microbes. Some heterotrophic microbes have evolved the ability to produce extracellular hydrolytic enzymes to catalyze the degradation of larger compounds (typically >600 Da) into smaller, more soluble substrates (Weiss *et al.*, 1991). These hydrolytic enzymes are secreted and function outside of the cell thereby allowing these heterotrophs to access and utilize more complex compounds. The activities and structural specificities of these hydrolytic enzymes influence the quantity and types of compounds that can be readily transformed and assimilated into biomass vs. the fractions that remain intact (e.g., Hoppe 1993, Meyer-Reil 1991). Thus, extracellular hydrolytic enzymes are often considered to be a rate-limiting step in the carbon remineralization process in marine environments (Arnosti, 2011). Similarly, the production of extracellular hydrolytic enzymes has been proposed to be a key microbial trait with respect to the decomposition of litter in soils (Schimel and Weintraub, 2003; Fontaine and Barot, 2005; Moorhead and Sinsabaugh, 2006; Allison *et al.*, 2010; Allison, 2012).

Our results suggest that extracellular hydrolytic enzymes may be a key trait when considering the lability and remineralization of natural organic matter in marine sediments. Moreover, gene dose (i.e., the number of gene copies for a given enzyme) has been suggested to play a key role in dictating the hydrolytic power of a strain (Enke *et al.*, 2018). In our study, we observed that greater hydrolytic power not only allowed *Pseudoalteromonas* sp. 3D05 to more readily access a protein-rich carbon pool, but also led to substantially more total organic matter being remineralized (higher total CO_2_ production by this strain). However, it may be possible that these observations are due to physiological differences in terms of cellular allocation and utilization of carbon substrates rather than solely hydrolytic power. Further analyses are needed to quantify these hydrolytic enzymes during our incubations in order to confirm their expression and activity. Nevertheless, our results support the notion that extracellular hydrolytic enzymes may be an important microbial trait when considering the quantities and types of organic matter that are accessed and consumed by heterotrophs.

Although the ability to produce and secrete hydrolytic enzymes is an ecologically important metabolic trait, the range of species that can produce these hydrolytic enzymes is narrow and often limited to a minor fraction of heterotrophs in complex communities (Langenheder *et al.*, 2006; Logue *et al.*, 2016). Consequently, the ability to produce hydrolytic enzymes is thought to be a functionally dissimilar trait, rather than functionally redundant (Rivett and Bell, 2018) This implies that differences in microbial community composition will result in variations in carbon remineralization rates independent of environmental conditions. For example, the addition of high molecular weight DOM to seawater mesocosms prepared from distinct water masses led to increased extracellular hydrolytic activity as well as growth of ‘rare’ OTUs (Balmonte *et al.*, 2019). Moreover, the spectrum of active enzymes detected in these mesocosms varied considerably, which suggested that the substrates that would have been consumed also differed among communities. The metabolic factors that lead to the expression and function of hydrolytic enzymes in marine environments are mostly matters of speculation. Studies suggest that inter-species interactions and cell numbers (Krupke *et al.*, 2016; Cezairliyan *et al.*, 2017) can influence the expression of these enzymes. Future work is needed to determine the precise mechanisms which control the production of hydrolytic enzymes in order to resolve the microbial controls on the marine carbon cycle.

## Acknowledgements

This work was supported by grants from the Gordon and Betty Moore Foundation (to A.P.) and by Center for Dark Energy Biosphere Investigations (to N.M.). We thank Susan Carter, Julia Schwartzman, and Maria Pachiadaki for analytical assistance as well as NOSAMS for isotopic measurements. Targeted on- and off-axis sampling in Guaymas Basin was made possible by NSF grant 1357238 to A.T., and the excellent performance of the *Alvin* and *Sentry* teams during cruise AT37-06.

